# Association between social factors and gastrointestinal parasite product excretion in a group of non-cooperatively breeding carrion crows

**DOI:** 10.1101/2020.03.20.999987

**Authors:** Claudia A.F. Wascher

## Abstract

The social environment strongly affects the physiological stress response in group living animals, which in turn can affect the immune system and increase susceptibility to parasites. Here, I investigate relationships between social behavior and gastrointestinal parasite product excretion in the carrion crow (*Corvus corone*). Individuals from a population of non-cooperatively breeding carrion crows excreted less samples containing coccidian oocysts when kept in larger groups (8 or 9 individuals) compared to those individuals kept in smaller groups (2 or 3 individuals). Lower-ranking individuals excreted more samples containing parasite oocysts compared to higher-ranking individuals. The strength of affiliative relationships and number of related individuals in the group did not relate to the proportion of droppings containing coccidian oocysts. The present results confirm an association between social environment and parasite excretion patterns in carrion crows, but the patterns described in the present study differ from previously published data derived from a group of cooperatively breeding crows. This illustrates that differences between the social systems of carrion crows might result in different associations between the social environment and parasite product excretion patterns.

**Significance statement:** One major cost of group living is an increase in susceptibility to parasites, however not all group living animals are affected by this in the same way. A better understanding how social behavior is associated with parasite burden can help to better understand evolution of group living. This study attempts to investigate associations between dominance rank, affiliative relationships as well as groups size and gastrointestinal parasite product excretion in a group of captive carrion crows. Lower-ranking individuals excreted more samples containing parasite oocysts compared to higher-ranking individuals, confirming an association between social relationships within the groups (for example dominance rank) and parasite excretion patterns.

## Introduction

Increased parasite burden is considered a major cost of group living (Alexander 1974). Initially group size has been hypothesized to be positively associated with parasite infection risk (Cote and Poulin 1995; Loehle 1995), whereas recent research hints towards the possibility of group size being a weak predictor of parasite infection risk (Rifkin et al. 2012) or negatively associated with parasite intensities (Patterson and Ruckstuhl 2013). Instead, behavior of animals in social groups for example frequency of social interactions and connectedness affects infection rates among group members (Rimbach et al. 2015; Balasubramaniam et al. 2019; Habig et al. 2019). Next to social interactions (Duboscq et al. 2016; Romano et al. 2016; VanderWaal et al. 2016), physiological processes that allow for increased parasite replication or survival in the host can significantly affect an individual’s susceptibility to parasites. For example, social behavior can significantly affect an individual’s physiological stress response (Wascher et al. 2009; Wittig et al. 2015), which can affect the immune system and make individuals more susceptible to parasite infections (Apanius 1998; Akinyi et al. 2019). Otherwise, parasitic infections affect an individual’s ability to engage in social behavior (Sheridan et al. 1994; von Holst 1998; Hanley and Stamps 2002; DeVries et al. 2003; Klein 2003; Lopes et al. 2016). Behavioral effects on an individual’s physiology and immune system range from relatively short-term, as in the effect on disease susceptibility (McEwen et al. 1997), to long-term, as in the effects on reproductive outcome (Buchholz 1995; Marzal et al. 2005; Hillegass et al. 2010) and have serious impacts on host longevity (Rousset et al. 1996; Archie et al. 2014).

The social environment can either facilitate or inhibit susceptibility and exposure to parasitism. Adverse effects of the social environment on health and susceptibility to parasites may be caused for example by increased competition and aggressive behavior (Azpiroz et al. 2003; Hawley et al. 2006; Chester et al. 2010). In meerkats, receiving but not initiating aggressive interactions was positively correlated with increased susceptibility of tuberculosis infection (Drewe 2010). Social status also affects an individual’s risk to be infected with parasites. A recent meta-analysis shows dominant individuals to suffer a higher risk to be infected compared to subordinate individuals and this effect to be mediated by social system (linear versus egalitarian hierarchies) and mating effort (Habig et al. 2018). In baboons (*Papio cynocephalus*), high-ranking males were less likely to become ill, and they recovered more quickly than low-ranking males (Archie et al. 2012).

Affiliative social interactions hold a risk of increased disease and parasite transmission. In rhesus macaques, *Macaca mulatta*, allo-grooming mediated transmission of *Escherichia coli* and central individuals in the social network could be considered ‘super-spreaders’ (Balasubramaniam et al. 2019). In Cape ground squirrels, *Xerus inauris*, increased durations of allo-grooming was associated with lower counts of ectoparasites (Hillegass et al. 2008). Affiliative behaviors also affect physiological processes which might affect the immune system, for example they have a stress-reducing effect (Sachser et al. 1998; Frigerio et al. 2003; Stöwe et al. 2008; Young et al. 2014; Müller-Klein et al. 2019) and might positively affect an individual’s immune system. In a captive population of cooperatively breeding carrion crows, individuals with strong affiliative relationships excreted less samples containing coccidian oocysts (Wascher et al. 2019).

In the present study, I investigate associations between social behavior and gastrointestinal parasite burden in the carrion crow, *Corvus coro*ne. Corvids express high variability in their social organization depending on life history and ecological factors. Within species the social organization might vary between different life history stages, seasons or populations. For example, in most European populations, carrion crows form socially monogamous pairs during the breeding season and large flocks during the rest of the year (Meide 1984; Glutz von Blotzheim 1985), whereas in northern Spain crows live in stable social groups of up to nine individuals, consisting of the breeding pair and retained offspring as well as immigrants, which are in most cases male individuals (Baglione et al. 2003). Group living in carrion crows is linked to a higher level of cooperation, for example in nestling provisioning (Baglione et al. 2005). Within their social groups, corvids establish valuable social relationships, characterized by spatial proximity, high levels of tolerance, relatively low frequencies of aggressive interactions and relatively high frequencies of affiliative behaviors (Fraser and Bugnyar 2010a; Heinrich 2011). Within such valuable relationships, individuals support another in agonistic encounters (Emery, et al. 2007; Fraser and Bugnyar 2012) and share information and resources (Bugnyar et al. 2001; de Kort et al. 2003). Affiliative relationships, are preferably formed with kin, but also non-kin individuals, in which case, long-term monogamous pair-bonds may ultimately develop (Loretto et al. 2012). Monogamous pair-bonds in corvids are characterized by complex social interactions, that demand a high level of cooperation, coordination and affiliation between the paired individuals, which may explain, in turn, the evolution of advanced cognitive skills (‘relationship intelligence’: Emery et al. 2007; Wascher et al. 2018). Social relationships, facilitate individuals in gaining access to resources, for example food or territories, and individuals in social relationships are more likely to win agonistic encounter through coalitional support (Bugnyar 2013). Outside such valuable social relationships corvids act more competitively (Bugnyar and Heinrich 2005; Bugnyar and Heinrich 2006) and usually establish linear dominance hierarchies (Izawa and Watanabe 2008). In a previous study on a cooperatively breeding population of captive carrion crows, strength of affiliative social relationships, and group size but not dominance rank or sex correlated with excretion of gastrointestinal parasite eggs and oocysts. Individuals with strong affiliative bonds excreted a smaller proportion of samples containing coccidian oocysts as did individuals living in larger groups (Wascher et al. 2019).

In the present study, I investigate the association between aspects of the social environment of non-cooperatively breeding captive carrion crows (strength of affiliative relationships, dominance hierarchy, and group structure) and fecal egg count. Fecal egg counts provide a reliable estimate of parasite infection rates (Seivwright et al. 2004; Daş et al. 2011). Further, I compare the results of the present study with the previously published data in Wascher et al. (2019). I expect a positive social environment, such as engaging in strong affiliative relationships, to reduce excretion of parasite eggs and oocysts. In the previous study, individual position in the dominance hierarchy was not correlated with eggs and oocysts excretion patterns. In the present study, I expect to confirm these results. Lastly, I investigate the association between group size (*i.e*. pairs or trios versus flocks of eight or nine individuals) and parasite product excretion. Parasite transmission may be expected to be facilitated by an increase in group size, however, in line with Wascher et al. (2019), I expect to find a negative association between group size and the number of parasite eggs and oocysts excreted by individuals.

## Methods

### Study subjects and ethics statement

I collected the data for this study in four phases, between 2008-2010 and between 2012-2015 from a population of captive carrion crows housed in large outdoor aviaries at the Konrad Lorenz research station (KLF), Grünau, Upper Austria (47°51’02 N 13°57’44 E). I observed 21 individuals (10 males and 11 females), kept in different group formations, such as groups of eight or nine individuals and pairs or trios. Due to the long-term character of the present study, individual birds were opportunistically moved between different compartments and group compositions due to age, reproduction, group expansion or the death of individual birds (Table 1). Aviaries were approximately 20-45 m^2^ and were equipped with wooden perches, natural vegetation and rocks. At the start of the study in 2008 groups have been kept in the large aviary (45 m^2^) and were consecutively moved into smaller compartments when separated into pairs and trios. In January 2012, all birds were moved into same sized compartments (20 m^2^). An enriched diet consisting of fruit, vegetables, bread, meat and milk products was provided on a daily basis. Water was available *ad libitum* for both drinking and bathing. This study complied with Austrian and local government guidelines. Individuals remain captivity housed in the Cumberland game park and the KLF (under the license AT00009917), before and after completion of the present study.

**Table 1:**
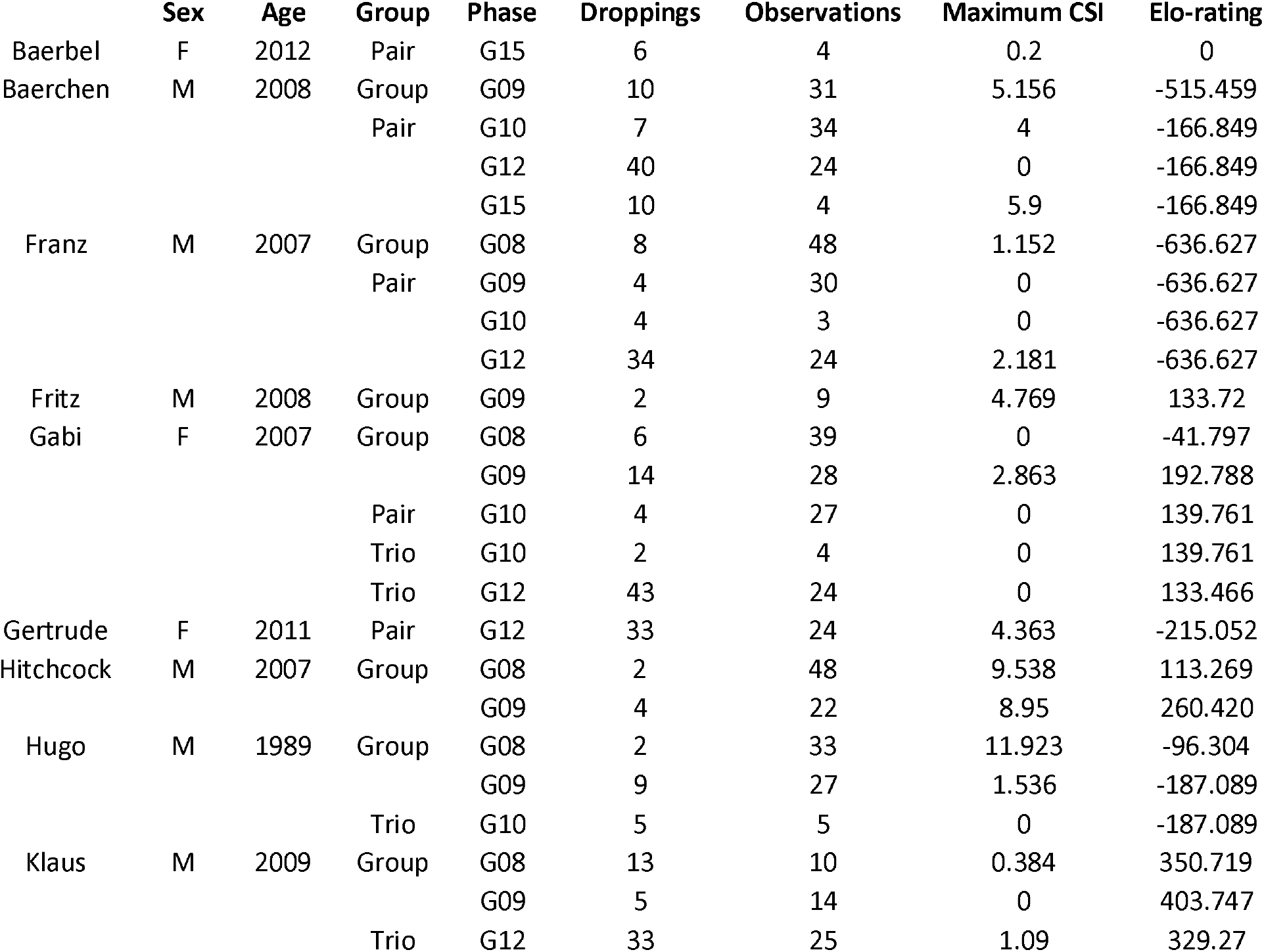
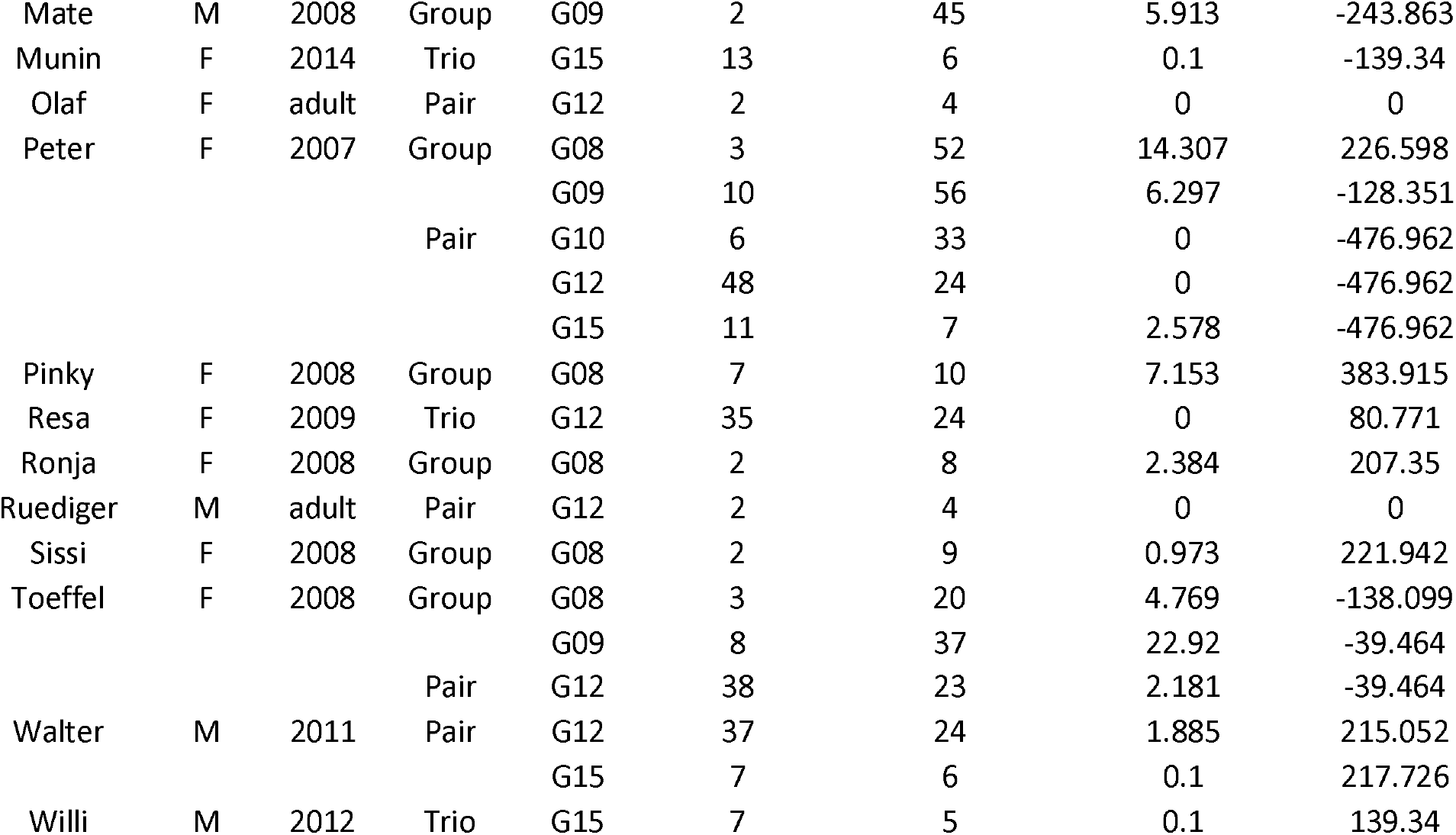
List of all focal individuals and information about population, sex (F = female, M = male), age (year of hatching; if not known, individuals are classified as adult), group (group composition: family, flock, pair, trio), phases of data taking during which the individual was recorded (G08: March to May, November to December 2008; G09: September to December 2009; G10: January to June 2010; G12: January to July 2012; G15: May to July 2015), number of droppings and behavioral focal observations collected, maximum composite sociality index (CSI) and Elo-rating.

### Behavioral data

I conducted a total of 899 individual focal observations. Each observation lasted five minutes, and I recorded all occurring behaviors. For this study, I focused on the frequencies of agonistic behavior (threat, chase flight, and fight) and affiliative behaviors (allopreen and contact sit). I recorded the identity, role (initiator/receiver) of interacting individuals and the outcome of the agonistic interaction (winner/loser), with the loser of an agonistic interaction defined as the individual that retreated.

### Composite sociality index

For each phase of data collection, I calculated a composite sociality index (CSI) for each crow dyad within a group according to Silk *et al.* (2010). I included two affiliative behaviors, namely contact sitting and allopreening, to calculate the CSI. The higher the CSI of a dyad, as compared with the frequency of the affiliative interactions observed within that dyad’s group, the stronger the affiliative bond between the two individuals in that dyad. For statistical analysis, I used the maximum CSI among all dyads for each individual, which reflected their strongest affiliative relationship within the group. For descriptive purposes, I classified dyads as ‘bonded’ when they displayed a higher CSI than the average of the entire sample and lower rates of aggression than the average of the entire group.

### Elo-rating

I calculated the relative success levels of individuals in agonistic encounters as an Elo-rating in the R package ‘*aniDom*’ (version 0.1.4; Sánchez-Tójar et al. 2018). Elo-rating allows to track dynamic changes in rank over the different phases of data collection. Each individual was rated based on the outcome of each discrete interaction (winner / loser) and the (predicted) probability of that outcome occurring (Neumann et al. 2011).

### Parasitological examination

During the entire study period, I collected a total of 559 individual droppings directly after defecation (for a detailed overview see table 1). I determined individual gastrointestinal parasite load from droppings. From 2008 until November 2011, I used a modified version of the flotation method (Schnieder et al. 2006). I suspended the fresh droppings (0.1 g) in a 2 ml collection tube with 1 ml saturated saline. I shook collection tubes for 10 seconds and afterwards centrifuged for 5 minutes at 3000 rpm. After centrifugation, I filled the collection tubes with saline solution and positioned a cover slip (18 × 18 mm) onto the tube. The high density of the saline solution causes the parasite eggs and oocysts to float up and be caught on the cover slip (Carta and Carta 2000). After 10 minutes, I moved the cover slip onto an object slide and identified and counted the parasite eggs and oocysts (by size and shape). From December 2011 onwards, I used a McMaster counting chamber. I weighted the entire dropping, then diluted with 3 ml saturated NaCl solution per 0.1 g of dropping and mixed thoroughly. Afterwards, I poured the solution into both compartments of the McMaster counting chambers. After a 10-minutes resting period, I counted the number of parasite eggs and oocysts in each compartment and calculated the number of parasite products per 1 ml of dropping.

I used a compound microscope with 100-fold and 400-fold amplification for parasite examination to identify coccidian oocysts, several nematode species (*Capillaria* sp., *Ascarida* sp., *Syngamus* sp. and *Heterakis* sp., *Trichostrongylus tenius*) and cestodes to a varying degree Table 2. I used presence versus absence of parasite eggs and oocysts for further analysis, which allowed a direct comparison between the two applied methods of droppings examination (flotation versus McMaster). As only nine samples contained cestodes, I conducted no further statistical analysis on this parasite group.

**Table 2.** Number of samples containing and not containing different parasite products.

|  | Number of samples<br>containing | Number of samples<br>not containing | Total number of<br>samples |
| --- | --- | --- | --- |
| nematode eggs | 387 | 172 | 559 |
| coccidian oocysts | 316 | 243 | 559 |
| cestodes | 9 | 550 | 559 |

### Data analysis

I analyzed factors affecting the proportion of droppings containing coccidian oocysts and nematode eggs in crows using the *glmer* function in R (version 3.5.3; R Core Team 2019) in the *lme4* package (version 1.1-19; Bates et al. 2015). In two models, the number of samples containing nematode eggs or coccidian oocysts for each period of data collection was the response term. I calculated GLMMs with binomial error distribution and a two-vector response variable comprising the number of infected and non-infected samples for each individual in each phase. I employed various model diagnostics to confirm model validity (visual inspection of the distribution of residuals, Q-Q plots, residuals plotted against fitted values), none of which suggested violation of the model’s assumptions. To assess multicollinearity between fixed factors, I calculated variance inflation factors (VIFs) using the *vif* function in the package *car* (version 3.0-6; Fox and Weisberg 2011). VIFs for all models were below 1.6, indicating that there was no issue with multicollinearity (Zuur et al. 2009). Strength of affiliative relationships (CSI value), social structure (pair/trio or group), number of related individuals, sex and Elo-rating were included as explanatory variables. For each model, I fitted individual identity as a random term to control for the potential dependence associated with multiple samples from the same individuals. The statistical significance level was adjusted to P ≤ 0.025 following Bonferroni, to account for multiple testing of coccidia oocysts and nematode eggs. In addition to the main analysis, I compared the results based on the present dataset (collected from a non-cooperatively breeding population of captive carrion crows) with those based on a previously published data (from a cooperatively breeding population) (Wascher, et al. 2019). Data collection and analysis (for example calculation of CSI and Elo-rating scores) was comparable in both studies. In order to investigate potential differences in social structure between the two populations, I compared the number of social relationships applying a GLMM with Poisson error distribution, as well as CSI value and Elo-rating applying two general linear models (GLMs) with Gaussian error distribution. Population included as explanatory variable, individual identity was fitted as a random term in each model and I calculated models in the *lme4* package.

## Results

### Social relationships

I observed 30 bonded dyads (out of 213 dyads in total), of which 19 were male-female dyads (five between related individuals and 14 between unrelated individuals). Six dyads were male-male dyads (all between unrelated individuals) and five female-female dyads (all between unrelated individuals). On average (± SD), males and females had 2.125 (± 1.642) and 1.625 (± 1.505) bonded partners, respectively. The mean CSI (± SD) between bonded dyads was 4.122 (± 2.612) for male-female, 2.561 (± 1.154) for female-female and 3.911 (± 2.062) for male-male bonds. Neither the number of social bonds, the CSI value or Elo-rating differed between the cooperatively breeding and non-cooperatively breeding populations of crows (number of social bonds: estimate ± SE = −0.085 ± 0.361, z = −0.235, P = 0.813; CSI: estimate ± SE = −0.582 ± 0.767, z = −0.758, P = 0.448; Elo-rating: estimate ± SE = 0.001 ± 0.001, z = 1.113, P = 0.265).

### Occurrence of coccidian oocysts and nematode eggs

243 samples from 18 individuals contained coccidian oocysts (43 % compared to 31 % in the Spanish population of cooperatively breeding crows). Crows kept in groups of eight or nine individuals excreted less samples containing coccidian oocysts compared to crows kept in pairs or trios (estimate ± SE = 0.986 ± 0.276, z = 3.564, P < 0.001, Fig. 1). Higher ranking individuals excreted fewer samples containing coccidian oocysts than lower ranking individuals (estimate ± SE = −0 ± 0, z = −2.167, P = 0.03, Fig. 2). However, the number of samples containing coccidian oocysts was not related to number of related individuals in the group, CSI (after Bonferroni correction) and sex (Table 3a).

**Table 3.** Results of the generalized mixed linear model investigating factors affecting patterns of coccidian oocyst and nematode egg excretion. Models investigate effects of group structure, number of related individuals (Nr related), strength of affiliative relationships (CSI), sex and dominance hierarchy (Elo-rating) on presence or absence of (a) coccidian oocysts and (b) nematode eggs in the sample. Significant values (*p*≤0.05) are highlighted in bold.

| | Parameters | Estimate ± SE | *z* | *p* |
| --- | --- | --- | --- | --- |
| (a) Coccidia | Intercept | **-1.28 ± 0.302** | **-4.229** | **<0.001** |
|  | Group structure | **0.986 ± 0.276** | **3.564** | **<0.001** |
| | Nr related | -1.134 ± 0.632 | -1.794 | 0.072 |
| | CSI | 0.072 ± 0.032 | 2.209 | 0.027 |
|  | Elo-rating | **-0 ± 0** | **-2.167** | **0.03** |
| | Sex | -0.007 ± 0.183 | -0.038 | 0.969 |
| (b) Nematodes | Intercept | **-1.181 ± 0.506** | **-2.333** | **<0.001** |
| | Group structure | -0.169 ± 0.318 | 0.53 | 0.595 |
| | Nr related | -0.964 ± 0.619 | -1.556 | 0.119 |
| | CSI | -0.001 ± 0.044 | -0.039 | 0.968 |
| | Elo-rating | 0 ± 0 | 0.053 | 0.957 |
| | Sex | 0.043 ± 0.587 | 0.073 | 0.941 |

**Figure 1.**
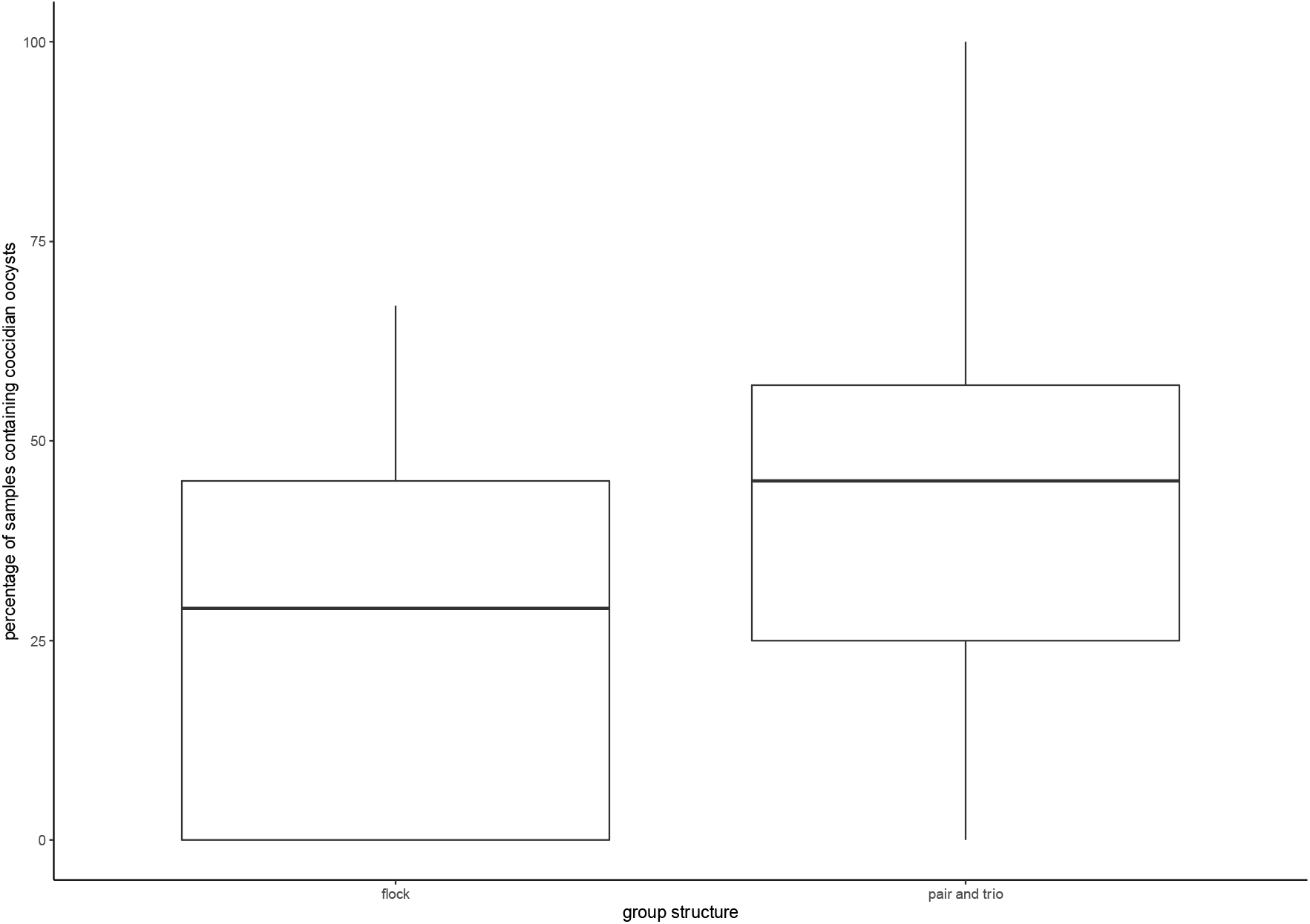
Percentage of samples containing coccidian oocysts in carrion crow droppings in relation to the group structure. Box plots show the median and the interquartile range from the 25th to the 75th percentiles.

**Figure 2.**
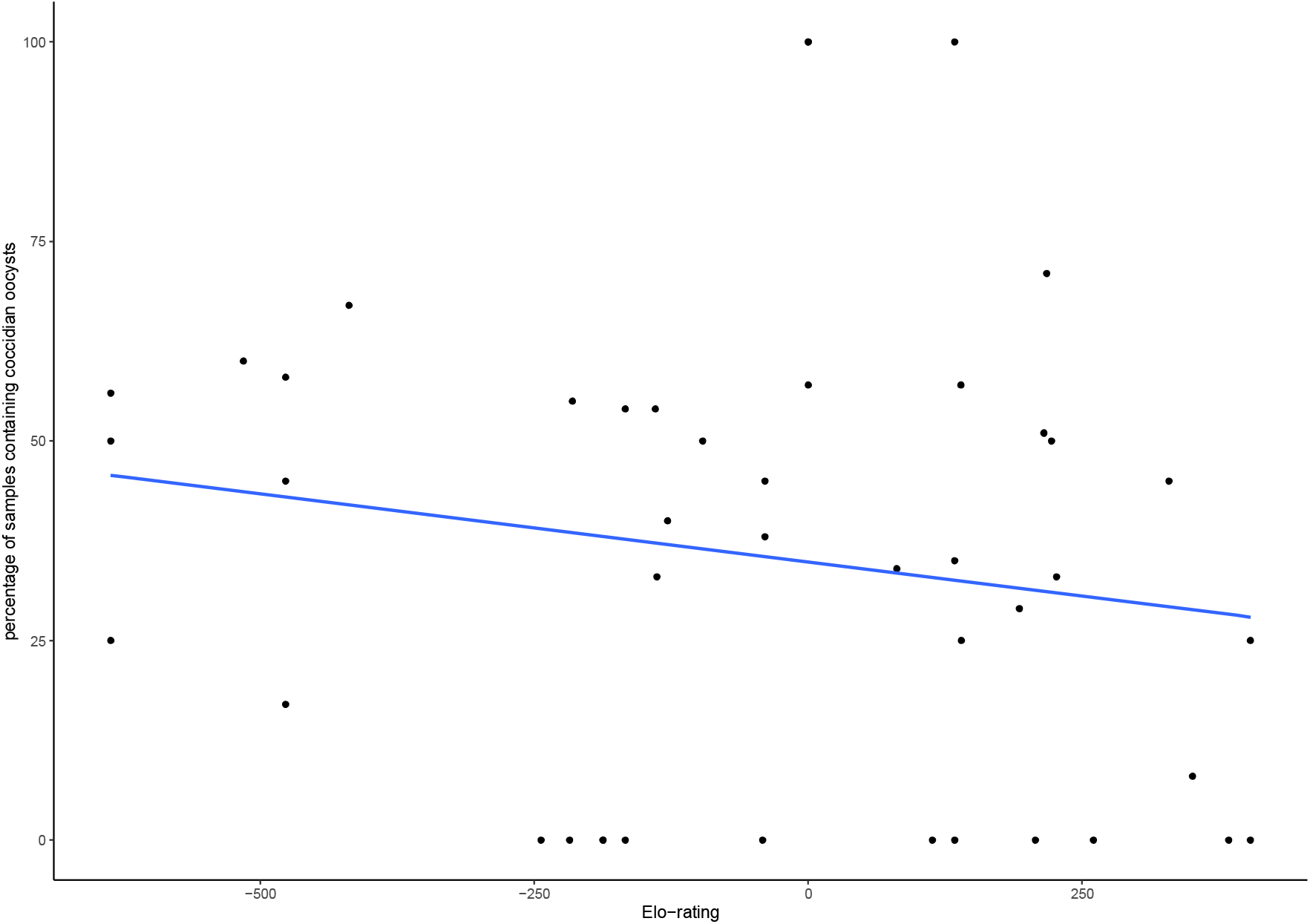
Percentage of samples containing coccidian oocysts in carrion crow droppings in relation to individual Elo-ratings.

Overall, 172 samples, collected from 14 individuals contained nematode eggs (31 % compared to 9 % in the Spanish population of cooperatively breeding crows). None of the factors investigated significantly affected excretion patterns of nematode eggs (Table 3b).

## Discussion

In this study, individuals from a population of non-cooperatively breeding carrion crows excreted less samples containing coccidian oocysts when kept in larger groups (8 or 9 individuals) compared to those individuals kept in smaller groups (2 or 3 individuals). Further, lower-ranking individuals excreted more samples containing parasite oocysts compared to higher-ranking individuals. A similar association between group-size and parasite excretion patterns was found in a population of cooperatively breeding carrion crows (Wascher et al. 2019). However, other patterns in the present study contrast did not replicate previous findings. In a group of cooperatively breeding crows, the proportion of samples containing coccidian oocysts was negatively correlated with the strength of affiliative social relationships, but dominance hierarchy did not affect parasite excretion patterns (Wascher et al. 2019).

The present study suggests that variation in the social system within a species can result in differences in parasite product excretion patterns. Here, in a population of non-cooperatively breeding carrion crows, subordinate individuals excreted more samples containing coccidian oocysts. In contrast, no such association was found in a population of cooperatively-breeding carrion crows (Wascher et al. 2019). In a way this is surprising, as dominance is a mechanism to suppress reproduction in subordinate individuals in some cooperatively breeding mammalian species (Creel et al. 1992; Young et al. 2006) and therefore I would have expected an effect of rank position onto parasite excretion patterns in the cooperatively breeding population of carrion crows, but not in the non-cooperatively breeding group. On the other hand, avian cooperatively breeding societies have often been described highly tolerant, especially towards related individuals (Baglione et al. 2003; Dickinson et al. 2009), which could explain the presented results. The relationship between social rank, glucocorticoid excretion and immune function is a complex one and can depend on factors like mating system, costs of rank acquisition and maintenance, or social stability (Goymann and Wingfield 2004; Cavigelli and Chaudhry 2012; Beehner and Bergman 2017; Habig et al. 2018). Further, the relationship between glucocorticoids and parasite burden might not be linear, and only high levels of physiological stress might influence parasite infection (Romeo et al. 2020). In two recent meta-analysis on several vertebrate taxa, dominant individuals exhibited higher parasite burden compared to subordinates, especially in linear versus egalitarian hierarchies and in mating systems where dominance rank predicts mating effort (Habig and Archie 2015; Habig et al. 2018). This is in contrast to the results of the present study. Carrion crows also can be considered to form linear and stable hierarchies (Chiarati et al. 2010), with dominant individuals gaining priority access to resources such as food (Chiarati et al. 2011). In Siberian hamsters, *Phodopus sungorus*, social defeat affects immune function (Chester et al. 2010). Similar to the results in the present study, subordinate individuals display a higher parasite burden in a variety of species (Guenons, Cercopithecus mitis: (Foerster et al. 2015). Higher levels of parasite excretion in subordinate individuals might reflect higher exposure to psychosocial stressors compared to dominant individuals (Levy et al. 2020).

I did not find an effect of sex on parasite excretion patterns. Differences in parasite burden between the sexes are mostly mediated by endocrine-immune interactions (Zuk and McKean 1996; Klein 2004). For example, high levels of testosterone are expected to be positively correlated with high parasite burden (Hudman et al. 2000; Decristophoris et al. 2007). Related to this, males are usually expected to have higher levels of endoparasites compared to females. However, this pattern does not hold in all species. Similar to the present study in carrion crows, a relationship between sex and parasite excretion was missing for example in red-fronted lemurs, *Eulemur fulvus rufus* (Clough et al. 2010), brown mouse lemur, *Microcebus rufus* (Rafalinirina et al. 2019), and four species of songbirds (Granthon and Williams 2017). Data for the present study was collected throughout the year in non-reproductively active birds, hence sex-steroid levels are expected to be generally low (Soma 2006; Hau et al. 2008), and therefore both sexes might have been similarly susceptibly to parasite infection in the studied population.

Increased exposure to parasites and disease transmission is considered as one of the major disadvantages of group living. Group size is usually positively related to parasite burden, which is seen as a major selective force in group living animals (Cote and Poulin 1995; Rifkin et al. 2012; Patterson and Ruckstuhl 2013). Similar to a previous study in captive cooperatively breeding carrion crows, individuals in the present study excreted less samples containing coccidian oocysts when living in larger groups. A limitation of the present study is the fact that I only used limited number of different group sizes (2-3 and 8-9 individuals). Individuals were grouped together in these categories opportunistically. Juvenile crows in the wild form ‘non-breeder flocks’ (Meide 1984; Glutz von Blotzheim 1985), usually larger than 8-9 individuals, however group sizes at the study site needed to be keep small in order for them to be manageable. As adults, individuals typically form male-female pairs, and this was the preferred social unit size I tried to achieve. Due to reasons of animal management, I temporarily had to keep crows as trios, when I did not have additional individuals available to form pairs. At all times I monitored groups of crows and in cases were levels of aggression increased between individuals, groups have been separated into smaller groups. It had to be considered that groups size and parasite load are associated in a non-linear manner, which could not be assessed in the present study (Hopkins et al. 2020). Infection by parasites in group living animals can be caused by different mechanisms. For example, parasites can be transmitted directly during social interactions, hence individuals engaging more socially are at greater risk to be infected. Parasite infection can also be related to physiological processes, which allow parasites increased replication or survival in the host, for example a compromised immune system, mediated by the physiological stress response (Muehlenbein and Watts 2010). Unfortunately, in the present study I have not measured individual stress levels and therefore cannot verify an association between parasite excretion patterns and levels of stress. The relationship between the physiological stress response and the immune system and related to this susceptibility to parasites is complex. A recent meta-analysis in 110 records from 65 studies in mammalian hosts from experimental and observational studies generally indicated a positive relationship between glucocorticoids and parasite burden (Defolie et al. 2020), however overall results about the relationship are complex. For example, in male chimpanzees, *Pan troglodytes schweinfurthii*, intestinal parasite prevalence and richness is positively associated with individual stress levels (Muehlenbein and Watts 2010). In chickens, *Gallus domesticus*, increased corticosterone levels were associated with increased numbers of coccidia oocysts and the length of the excretion period (Graat 1996) and in baboons, *Papio cynocephalus*, infection with helminths was associated with higher glucocorticoid levels (Akinyi et al. 2019). Other studies report a lack of correlation between physiological measures of stress and parasite burden, for example in male black grouse, *Tetrao tetrix* (Sokół and Koziatek-Sadłowska 2020) and Blue-crowned manakins, *Lepidothrix coronata*, (Bosholn et al. 2020) individual corticosterone levels were not correlated with parasite infection. Future studies investigating the direct link between social behavior, physiological stress response and parasite excretion patterns would be desirable.

In contrast to (Wascher et al. 2019), strengths of affiliative social relationships was not associated with the proportion of samples containing coccidian oocysts. Neither the strength nor the number of affiliative relationships differed significantly between the subject populations in these studies. Therefore, it does not seem that populations show a general, behavioral difference in affiliative relationships within groups. In cooperatively breeding crows, individuals with strong affiliative relationships excreted less samples containing coccidian oocysts (Wascher et al. 2019), which could reflect a stress reducing and immune system enhancing function of affiliative social relationships. Strong relationships between pair-partners, between kin, and also between non-paired and unrelated individuals play a significant role in the social life of corvids, and have been hypothesized to be an important driver in the evolution of cognition (Emery et al. 2007). Social relationships in common ravens have been shown to facilitate access to resources (Braun and Bugnyar 2012) and cooperation (Fraser and Bugnyar 2012). Following conflicts, common ravens reconcile and consolidate valuable relationships, therefore mitigating potential costs of disrupted relationships (Fraser and Bugnyar 2010b; Fraser and Bugnyar 2011). Based on these previous findings, it can be assumed that strong affiliative relationships hold significant benefits for corvids. Therefore, I expected to find, but was unable to confirm in the present study, that the nature of an individual’s social relationships would affect their health and physiology. This is especially interesting as dominance rank was negatively correlated with the proportion of samples excreted containing coccidian oocysts. Aggressive social interactions have previously been shown to be among the most potent stressors (Wascher et al. 2008) and position in the hierarchy is related to stress (Creel 2001) and immunodepression (Barnard et al. 1998) in a number of species. Social relationships have been suggested to mitigate the effects of aggression onto the physiological stress response (Sachser et al. 1998) and therefore I would have expected to find an effect of strength of affiliative relationships on parasite product excretion in carrion crows.

In the present study, the excretion of coccidian oocysts but not excretion of nematode eggs was found to be related to social factors. These results replicated those of Wascher et al. (2019) and may be related to differences between these parasite species in their lifecycles. Coccidian oocysts have a shorter prepatent period compared to nematode eggs (Edgar 1955; French and Zachary 1994). This could potentially, make them more sensitive to short-term changes in stress levels and immune system. This is supported by a study in carrion crows, showing coccidian oocysts but not nematode eggs to significantly increase in the first week after a major stressor (Spreafico et al. 2012) and in graylag geese excretion of coccidian oocysts but not nematode eggs are significantly increased in the first week after social isolation (Ludwig et al. 2017).

In summary, more droppings containing coccidian oocysts were excreted by individuals kept in pairs and trios compared to groups of eight or nine individuals. Strength of affiliative social relationships did not correlate with parasite product excretion; however subordinate individuals excreted more samples containing coccidian oocysts compared to dominant individuals. The present results illustrate differences in the social system in carrion crows also resulting in different associations between social environment and parasite product excretion.

## Acknowledgments

I am very grateful to Gaius de Smidt for detailed feedback on the manuscript. I also thank two anonymous referees for valuable comments on the manuscript.

## Ethics Statement

This work was supported by the Fonds zur Förderung Wissenschaftlicher Forschung Austria (FWF) project P21489-B17 to Kurt Kotrschal and CAFW, and permanent support was provided by the ‘Verein der Förderer’ and the Herzog von Cumberland Stiftung. The author declares no conflict of interest. All procedures were conducted in accordance with the ASAB/ ABS guidelines for the treatment of animals in behavioral research. The keeping of these captive birds was authorized under a license issued to the Cumberland Wildlife park Grünau (AT00009917).

## Notes

### Competing Interest Statement

The authors have declared no competing interest.

## References

Akinyi MY, Jansen D, Habig B, Gesquiere LR, Alberts SC, Archie EA. 2019. Costs and drivers of helminth parasite infection in wild female baboons. J Anim Ecol. 88(7):1029–1043.

Alexander RD. 1974. The evolution of social behavior. Annu Rev Ecol Evol Syst. 5:325–383.

Apanius V. 1998. Stress and immune defense. In: Advances in the Study of Behavior. Vol. 27. Elsevier. p. 133–153.

Archie EA, Altmann J, Alberts SC. 2012. Social status predicts wound healing in wild baboons. PNAS. 109(23):9017–9022.

Archie EA, Tung J, Clark M, Altmann J, Alberts SC. 2014. Social affiliation matters: both same-sex and opposite-sex relationships predict survival in wild female baboons. Proc R Soc B. 281(1793):20141261.

Azpiroz A, Garmendia L, Fano E, Sanchez-Martin JR. 2003. Relations between aggressive behavior, immune activity, and disease susceptibility. Aggress Violent Behav. 8(4):433–453.

Baglione V, Canestrari D, Marcos JM, Ekman J. 2003. Kin selection in cooperative alliances of carrion crows. Science. 300(5627):1947–9.

Baglione V, Marcos JM, Canestrari D, Griesser M, Andreotti G, Bardini C, Bogliani G. 2005. Does year-round territoriality rather than habitat saturation explain delayed natal dispersal and cooperative breeding in the carrion crow? J Anim Ecol. 74(5):842–851.

Balasubramaniam KN, Beisner BA, Hubbard JA, Vandeleest JJ, Atwill ER, McCowan B. 2019. Affiliation and disease risk: social networks mediate gut microbial transmission among rhesus macaques. Anim Behav. 151:131–143.

Barnard CJ, Behnke JM, Gage AR, Brown H, Smithurst PR. 1998. The role of parasite– induced immunodepression, rank and social environment in the modulation of behaviour and hormone concentration in male laboratory mice (Mus musculus). Proc R Soc Lond B. 265(1397):693–701.

Bates D, Mächler M, Bolker B, Walker S. 2015. Fitting linear mixed-effects models using lme4. J Stat Softw. 67(1):1–48.

Beehner JC, Bergman TJ. 2017. The next step for stress research in primates: To identify relationships between glucocorticoid secretion and fitness. Horm Behav. 91:68–83.

Bosholn M, Anciães M, Gil D, Weckstein JD, Dispoto JH, Fecchio A. 2020. Individual variation in feather corticosterone levels and its influence on haemosporidian infection in a Neotropical bird. Ibis. 162(1):215–226.

Braun A, Bugnyar T. 2012. Social bonds and rank acquisition in raven nonbreeder aggregations. Anim Behav. 84(6):1507–1515.

Buchholz R. 1995. Female choice, parasite load and male ornamentation in wild turkeys. Anim Behav.50(4):929–943.

Bugnyar T. 2013. Social cognition in ravens. CCBR. 8:1–12.

Bugnyar T, Heinrich B. 2005. Ravens, Corvus corax, differentiate between knowledgeable and ignorant competitors. Proc R Soc B. 272(1573):1641–1646.

Bugnyar T, Heinrich B. 2006. Pilfering ravens, Corvus corax, adjust their behaviour to social context and identity of competitors. Anim Cogn. 9(4):369–376.

Bugnyar T, Kijne M, Kotrschal K. 2001. Food calling in ravens: are yells referential signals? Anim Behav. 61(5):949–958.

Carta LK, Carta DG. 2000. Nematode specific gravity profiles and applications to flotation extraction and taxonomy. Nematology. 2(2):201–210.

Cavigelli SA, Chaudhry HS. 2012. Social status, glucocorticoids, immune function, and health: Can animal studies help us understand human socioeconomic-status-related health disparities? Horm Behav. 62(3):295–313.

Chester Emily M, Bonu T, Demas GE. 2010. Social defeat differentially affects immune responses in Siberian hamsters (Phodopus sungorus). Physiol Behav. 101(1):53–58.

Chester Emily M., Bonu T, Demas GE. 2010. Social defeat differentially affects immune responses in Siberian hamsters (Phodopus sungorus). Physiol Behav. 101(1):53–58..

Chiarati E, Canestrari D, Vera R, Marcos JM, Baglione V. 2010. Linear and stable dominance hierarchies in cooperative carrion crows. Ethology. 116(4):346–356.

Chiarati E, Canestrari D, Vila M, Vera R, Baglione V. 2011. Nepotistic access to food resources in cooperatively breeding carrion crows. Behav Ecol Sociobiol. 65(9):1791–1800.

Clough D, Heistermann M, Kappeler PM. 2010. Host intrinsic determinants and potential consequences of parasite infection in free-ranging red-fronted lemurs (Eulemur fulvus rufus). Am J Phys Anthropol. 142(3):441–452.

Cote IM, Poulin R. 1995. Parasitism and group size in social animals: a meta-analysis. Behav Ecol. 6(2):159–165.

Creel S. 2001. Social dominance and stress hormones. Trends Ecol Evol. 16(9):491–497.

Creel S, Creel N, Wildt DE, Monfort SL. 1992. Behavioural and endocrine mechanisms of reproductive suppression in Serengeti dwarf mongooses. Anim Behav. 43(2):231–245.

Daş G, Savaş T, Kaufmann F, Idris A, Abel H, Gauly M. 2011. Precision, repeatability and representative ability of faecal egg counts in Heterakis gallinarum infected chickens. Vet Parasitol. 183(1-2):87–94.

Decristophoris PMA, von Hardenberg A, McElligott AG. 2007. Testosterone is positively related to the output of nematode eggs in male Alpine ibex (Capra ibex) faeces. Evol Ecol Res.(9):1277–1292.

Defolie C, Merkling T, Fichtel C. 2020. Patterns and variation in the mammal parasite– glucocorticoid relationship. Biol Rev. 95(1):74–93.

DeVries AC, Glasper ER, Detillion CE. 2003. Social modulation of stress responses. Physiol Behav. 79(3):399–407.

Dickinson JL, Euaparadorn M, Greenwald K, Mitra C, Shizuka D. 2009. Cooperation and competition: nepotistic tolerance and intrasexual aggression in western bluebird winter groups. Anim Behav. 77(4):867–872. doi:10.1016/j.anbehav.2008.11.026.

Drewe JA 2010. Who infects whom? Social networks and tuberculosis transmission in wild meerkats. Proc R Soc B. 277(1681):633–642.

Duboscq J, Romano V, Sueur C, MacIntosh AJJ. 2016. Network centrality and seasonality interact to predict lice load in a social primate. Sci Rep. 6(1):22095.

Edgar SA. 1955. Sporulation of oocysts at specific temperatures and notes on the prepatent period of several species of avian coccidia. J Parasitol. 41(2):214.

Emery NJ, Seed AM, von Bayern Auguste MP, Clayton NS. 2007. Cognitive adaptations of social bonding in birds. Phil Trans R Soc B. 362(1480):489–505.

Emery NJ, Seed AM, von Bayern Auguste M.P, Clayton NS. 2007. Cognitive adaptations of social bonding in birds. Phil Trans R Soc B. 362(1480):489–505.

Foerster S, Kithome K, Cords M, Monfort SL. 2015. Social status and helminth infections in female forest guenons (*Cercopithecus mitis*): Rank and nematode infections in a forest guenon. Am J Phys Anthropol. 158(1):55–66.

Fox J, Weisberg S. 2011. An {R} Companion to Applied Regression. second. California: Sage Publications.

Fraser ON, Bugnyar T. 2010a. The quality of social relationships in ravens. Anim Behav. 79(4):927–933.

Fraser ON, Bugnyar T. 2010b. Do ravens show consolation? Responses to distressed others. Brosnan SF, editor. PLoS ONE. 5(5):e10605.

Fraser ON, Bugnyar T. 2011. Ravens reconcile after aggressive conflicts with valuable partners. Iwaniuk A, editor. PLoS ONE. 6(3):e18118.

Fraser ON, Bugnyar T. 2012. Reciprocity of agonistic support in ravens. Anim Behav. 83(1):171–177.

French RA, Zachary JF. 1994. Parasitology and pathogenesis of Geopetitia aspiculata (Nematoda: Spirurida) in zebra finches (Taeniopygia guttata): Experimental infection and new host records. J Zoo Wildlife Med.:403–422.

Frigerio D, Weiss B, Dittami J, Kotrschal K. 2003. Social allies modulate corticosterone excretion and increase success in agonistic interactions in juvenile hand-raised graylag geese (Anser anser). Can J Zool. 81:1746–1754.

Glutz von Blotzheim UN. 1985. Handbuch der Vögel Mitteleuropas. Glutz von Blotzheim UN, editor. Wiesbaden: Aula-Verlag.

Goymann W, Wingfield JC. 2004. Allostatic load, social status and stress hormones: the costs of social status matter. Anim Behav. 67(3):591–602.

Graat L. 1996. Epidemiology of Eimeria acervulina infections in broilers: an integrated approach. Wageningen: Wageningen Institute of Animal Science, Wageningen Agricultural University.

Granthon C, Williams DA. 2017. Avian malaria, body condition, and blood parameters in four species of songbirds. Wilson J Ornithol. 129(3):492–508.

Habig B, Archie EA. 2015. Social status, immune response and parasitism in males: a meta-analysis. Phil Trans R Soc B. 370(1669):20140109.

Habig B, Doellman MM, Woods K, Olansen J, Archie EA. 2018. Social status and parasitism in male and female vertebrates: a meta-analysis. Sci Rep. 8(1):3629.

Habig B, Jansen DAWAM, Akinyi MY, Gesquiere LR, Alberts SC, Archie EA. 2019. Multi-scale predictors of parasite risk in wild male savanna baboons (Papio cynocephalus). Behav Ecol Sociobiol. 73(10):134.

Hanley KA, Stamps JA. 2002. Does corticosterone mediate bidirectional interactions between social behaviour and blood parasites in the juvenile black iguana, Ctenosaura similis? Anim Behav. 63(2):311–322.

Hau M, Gill SA, Goymann W. 2008. Tropical field endocrinology: Ecology and evolution of testosterone concentrations in male birds. Gen Comp Endocrinol.157(3):241–248.

Hawley DM, Lindström K, Wikelski M. 2006. Experimentally increased social competition compromises humoral immune responses in house finches. Horm Behav. 49(4):417–424.

Heinrich B. 2011. Conflict, cooperation, and cognition in the common raven. Adv Stud Behav. 43:189–237.

Hillegass M a., Waterman JM, Roth JD. 2010. Parasite removal increases reproductive success in a social African ground squirrel. Behav Ecol. 21(4):696–700.

Hillegass MA, Waterman JM, Roth JD. 2008. The influence of sex and sociality on parasite loads in an African ground squirrel. Behav Ecol. 19(5):1006–1011.

von Holst D. 1998. The concept of stress and its relevance for animal behavior. In: Advances in the Study of Behavior. Vol. 27. Academic Press. p. 1–131.

Hopkins SR, Fleming-Davies AE, Belden LK, Wojdak JM. 2020. Systematic review of modelling assumptions and empirical evidence: Does parasite transmission increase nonlinearly with host density? Golding N, editor. Methods Ecol Evol. 11(4):476–486.

Hudman SP, Ketterson ED, Nolan V. 2000. Effects of time of sampling on oocyst detection and effects of age and experimentally elevated testosterone on prevalence of coccidia in male dark-eyed juncos. Auk. 117:1048–1051.

Izawa E-I, Watanabe S. 2008. Formation of linear dominance relationship in captive jungle crows (Corvus macrorhynchos): Implications for individual recognition. Behav Proc. 78(1):44–52.

Klein SL. 2003. Parasite manipulation of the proximate mechanisms that mediate social behavior in vertebrates. Phys Behav. 79(3):441–449.

Klein SL. 2004. Hormonal and immunological mechanisms mediating sex differences in parasite infection. Parasite Immunol. 26(6-7):247–264. d

de Kort SR, Emery NJ, Clayton NS. 2003. Food offering in jackdaws (Corvus monedula). Naturwissenschaften. 90(5):238–240.

Levy EJ, Gesquiere LR, McLean E, Franz M, Warutere JK, Sayialel SN, Mututua RS, Wango TL, Oudu VK, Altmann J, et al. 2020. Higher dominance rank is associated with lower glucocorticoids in wild female baboons: A rank metric comparison. Horm Behav. 125:104826.

Loehle C. 1995. Social barriers to pathogen transmission in wild animal populations. Ecology. 76(2):326–335.

Lopes PC, Block P, König B. 2016. Infection-induced behavioural changes reduce connectivity and the potential for disease spread in wild mice contact networks. Sci Rep. 6(1):31790.

Loretto M-C, Fraser ON, Bugnyar T. 2012. Ontogeny of social relations and coalition formation in common ravens (Corvus corax). Int J Comp Psych. 25:180–194.

Ludwig SC, Kapetanopoulos K, Kotrschal K, Wascher CAF. 2017. Effects of mate separation in female and social isolation in male free-living Greylag geese on behavioural and physiological measures. Behav Proc. 138:134–141.

Marzal A, de Lope F, Navarro C, Møller AP. 2005. Malarial parasites decrease reproductive success: an experimental study in a passerine bird. Oecologia. 142(4):541–545.

McEwen BS, Biron C a, Brunson KW, Bulloch K, Chambers WH, Dhabhar FS, Goldfarb RH, Kitson RP, Miller a H, Spencer RL, et al. 1997. The role of adrenocorticoids as modulators of immune function in health and disease: neural, endocrine and immune interactions. Brain Res Rev. 23:79–133.

Meide M. 1984. Raben- und Nebelkrähe. Magdeburg: Westarp Wissenschaften.

Muehlenbein MP, Watts DP. 2010. The costs of dominance: testosterone, cortisol and intestinal parasites in wild male chimpanzees. BioPsychoSocial Med. 4(1):21.

Müller-Klein N, Heistermann M, Strube C, Franz M, Schülke O, Ostner J. 2019. Exposure and susceptibility drive reinfection with gastrointestinal parasites in a social primate. Funct Ecol. doi:10.1111/1365-2435.13313.

Neumann C, Duboscq J, Dubuc C, Ginting A, Irwan AM, Agil M, Widdig A, Engelhardt A. 2011. Assessing dominance hierarchies: validation and advantages of progressive evaluation with Elo-rating. Anim Behav. 82(4):911–921.

Patterson JEH, Ruckstuhl KE. 2013. Parasite infection and host group size: a meta-analytical review. Parasitology. 140(7):803–813.

R Core Team. 2019. R: a language and environment for statistical computing. Vienna. http://www.r723project.org/.

Rafalinirina AH, Randrianasy J, Wright PC, Ratsimbazafy J. 2019. Effect of socio-ecological factors and parasite infection on body condition of Brown Mouse Lemur Microcebus rufus (Mammalia: Primates: Cheirogaleidae). J Threat Taxa. 11(6):13632–13643.

Rifkin JL, Nunn CL, Garamszegi LZ. 2012. Do animals living in larger groups experience greater parasitism? A meta-analysis. Am Nat. 180(1):70–82.

Rimbach R, Bisanzio D, Galvis N, Link A, Di Fiore A, Gillespie TR. 2015. Brown spider monkeys (*Ateles hybridus*): a model for differentiating the role of social networks and physical contact on parasite transmission dynamics. Phil Trans R Soc B. 370(1669):20140110.

Romano V, Duboscq J, Sarabian C, Thomas E, Sueur C, MacIntosh AJJ. 2016. Modeling infection transmission in primate networks to predict centrality-based risk: Individual Centrality and Infection Flow. Am J Primatol. 78(7):767–779.

Romeo C, Wauters LA, Santicchia F, Dantzer B, Palme R, Martinoli A, Ferrari N. 2020. Complex relationships between physiological stress and endoparasite infections in natural populations. Ferkin M, editor. Curr Zool.:1–9.

Rousset F, Thomas F, Meeûs T De, Renaud F. 1996. Inference of parasite-induced host mortality from distributions of parasitic loads. Ecology. 77(7):2203–2211.

Sachser N, Dürschlag M, Hirzel D. 1998. Social relationships and the management of stress. Psychoneuroendocrinology. 23(8):891–904.

Sánchez-Tójar A, Schroeder J, Farine DR. 2018. A practical guide for inferring reliable dominance hierarchies and estimating their uncertainty. J Anim Ecol. 87(3):594–608.

Schnieder T, Boch J, Supperer R. 2006. Veterinärmedizinische Parasitologie. 6th ed. Schnieder T, Boch J, Supperer R, editors. Berlin: Parey.

Seivwright LJ, Redpath SM, Mougeot F, Watt L, Hudson PJ. 2004. Faecal egg counts provide a reliable measure of *Trichostrongylus tenuis* intensities in free-living red grouse *Lagopus lagopus scoticus*. J Helminthol. 78(1):69–76.

Sheridan JF, Dobbs C, Brown D, Zwilling B. 1994. Psychoneuroimmunology: stress effects on pathogenesis and immunity during infection. Clin Microbiol Rev. 7(2):200–212.

Silk JB, Beehner JC, Bergman TJ, Crockford C, Engh AL, Moscovice LR, Wittig RM, Seyfarth RM, Cheney DL. 2010. Female chacma baboons form strong, equitable, and enduring social bonds. Behav Ecol Sociobiol. 64(11):1733–1747.

Sokół R, Koziatek-Sadłowska S. 2020. Changes in the corticosterone level in tooting male black grouse (Tetrao tetrix) infected with Eimeria spp. Poult Sci J. 99(3):1306–1310.

Soma KK. 2006. Testosterone and aggression: Berthold, birds and beyond. J. Neuroendocrinol. 18(7):543–551.

Spreafico M, Szipl G, Kotrschal K, Wascher CAF. 2012. Physiological and behavioural response in carrion crows (Corvus corone corone) after relocation to a new environment. Wien Tierärztl Monat. 99(Suppl 1):61.

Stöwe M, Bugnyar T, Schloegl C, Heinrich B, Kotrschal K, Möstl E. 2008. Corticosterone excretion patterns and affiliative behavior over development in ravens (Corvus corax). Horm Behav. 53(1):208–216.

Ungerfeld R, Correa O. 2007. Social dominance of female dairy goats influences the dynamics of gastrointestinal parasite eggs. Appl Anim Behav Sci. 105(1–3):249–253.

VanderWaal KL, Obanda V, Omondi GP, McCowan B, Wang H, Fushing H, Isbell LA. 2016. The “strength of weak ties” and helminth parasitism in giraffe social networks. Behav Ecol. 27(4):1190–1197.

Wascher CAF, Arnold W, Kotrschal K. 2008. Heart rate modulation by social contexts in greylag geese (Anser anser). J Comp Psych. 122(1):100–107.

Wascher CAF, Canestrari D, Baglione V. 2019. Affiliative social relationships and coccidian oocyst excretion in a cooperatively breeding bird species. Anim Behav. 158:121–130..

Wascher CAF, Kulahci IG, Langley EJG, Shaw RC. 2018. How does cognition shape social relationships? Phil Trans R Soc B. 373(1756):20170293.

Wascher CAF, Scheiber IBR, Weiß BM, Kotrschal K. 2009. Heart rate responses to agonistic encounters in greylag geese, Anser anser. Anim Behav. 77(4):955–961.

Wittig RM, Crockford C, Weltring A, Deschner T, Zuberbühler K. 2015. Single aggressive interactions increase urinary glucocorticoid levels in wild male chimpanzees. Siegel A, editor. PLoS ONE. 10(2):e0118695.

Young AJ, Carlson AA, Monfort SL, Russell AF, Bennett NC, Clutton-Brock T. 2006. Stress and the suppression of subordinate reproduction in cooperatively breeding meerkats. PNAS. 103(32):12005–12010.

Young C, Majolo B, Schülke O, Ostner J. 2014. Male social bonds and rank predict supporter selection in cooperative aggression in wild Barbary macaques. Anim Behav. 95:23–32.

Zuk M, Kim T, Robinson S, Johnsen T. 1998. Parasites influence social rank and morphology, but not mate choice, in female red junglefowl, Gallus gallus. Anim Behav.

Zuk M, McKean KA. 1996. Sex differences in parasite infections: Patterns and processes. Int J Parasitol. 26:1009–1024.

Zuur AF, Ieno EN, Walker NJ, Saveliev AA, Smith GM. 2009. Mixed Effects Models and Extension in Ecology With R. New York: Springer.

